# *hkb* is required for *DIP-α* expression and target recognition in the *Drosophila* neuromuscular circuit

**DOI:** 10.1101/2023.10.15.562341

**Authors:** Yupu Wang, Rio Salazar, Luciano Simonetta, Violet Sorrentino, Terrence J. Gatton, Bill Wu, Christopher G. Vecsey, Robert A. Carrillo

## Abstract

Our nervous system contains billions of neurons that form precise connections with each other through interactions between cell surface proteins (CSPs). In *Drosophila*, the Dpr and DIP immunoglobulin protein subfamilies form homophilic or heterophilic interactions to instruct synaptic connectivity, synaptic growth and cell survival. However, the upstream regulation and downstream signaling mechanisms of Dprs and DIPs are not clear. In the *Drosophila* larval neuromuscular system, *DIP-α* is expressed in the dorsal and ventral type-Is motor neurons (MNs). We conducted an F1 dominant modifier genetic screen to identify regulators of Dprs and DIPs. We found that the transcription factor, *huckebein* (*hkb*), genetically interacts with *DIP-α* and is important for target recognition specifically in the dorsal Is MN, but not the ventral Is MN. Loss of *hkb* led to complete removal of *DIP-α* expression. We then confirmed that this specificity is through the dorsal Is MN specific transcription factor, *even-skipped* (*eve*), which acts downstream of *hkb*. Genetic interaction between *hkb* and *eve* revealed that they act in the same pathway to regulate dorsal Is MN connectivity. Our study provides insight into the transcriptional regulation of *DIP-α* and suggests that distinct regulatory mechanisms exist for the same CSP in different neurons.

## Introduction

The way animals perceive and respond to the environment relies on precise and robust neuronal connections. During development, each neuron must identify the correct synaptic partners among thousands of potential targets. A prevalent model for instructing synaptic recognition, repulsion, and self-avoidance is through the interaction between unique cell surface proteins (CSPs). A major subset of CSPs belong to the immunoglobulin superfamily (IgSF) and play important roles in synaptic development and maintenance in both vertebrates and invertebrates. In the well-studied vertebrate retina, retinal ganglion cells require Dscams and Sidekicks (Sdks) 1 and 2 to avoid self-synapses and form stereotyped connections, respectively (Garrett et al., 2018; Krishnaswamy et al., 2015; Yamagata and Sanes, 2019). In hard-wired invertebrate nervous systems, such as *C. elegans*, the heterophilic interaction between two IgSF proteins, Syg1 and Syg2, is required for HSNL motor neuron synapse formation (Shen and Bargmann, 2003; Shen et al., 2004). In the *Drosophila* mushroom body, neurons utilize different isoforms of Dscam1 to discriminate self/non-self (Hattori et al., 2009; Wang et al., 2004; Zhan et al., 2004). Several IgSF CSPs have also been implicated in synaptic connectivity in the *Drosophila* larval neuromuscular system where two type-Is motor neurons (Is MNs) and ∼29 type-Ib motor neurons (Ib MNs) form stereotyped connection with 30 muscles in each hemisegment (Hoang and Chiba, 2001; Lnenicka and Keshishian, 2000; Wang et al., 2022). For example, the immunoglobulin proteins Fasciclin 2 (Davis et al., 1997; Winberg et al., 1998) and Fasciclin 3 (Chiba et al., 1995; Kose et al., 1997) are required for specific larval MNs to recognize their muscle targets.

Recent biochemical studies revealed two *Drosophila* immunoglobulin protein subfamilies, the Defective proboscis response proteins (Dprs, 21 members) and Dpr-interacting proteins (DIPs, 11 members) families, that form homophilic or heterophilic interactions to instruct synaptic connectivity, synaptic growth, and cell survival (Ashley et al., 2019; Barish et al., 2018; Bornstein et al., 2021; Brovero et al., 2021; Carrillo et al., 2015; Cheng et al., 2019; Cosmanescu et al., 2018; Özkan et al., 2013; Sergeeva et al., 2020; Tan et al., 2015; Venkatasubramanian et al., 2019; Xu et al., 2018; Xu et al., 2019; Xu et al., 2022). For example, the well-studied Dpr10-DIP-α interaction is necessary for innervation of the dorsal Is MN on muscle 4 (m4) as loss of either *dpr10* or *DIP-α* leads to complete loss of m4-Is innervation (Ashley et al., 2019). In addition, loss of *dpr10* or *DIP-α* in the optic lobe causes significant mistargeting and cell death of Dm12 medulla neurons, suggesting multifaceted roles for Dpr10-DIP-α interactions (Xu et al., 2018; Xu et al., 2022). Similarly, the recognition between yellow R7 photoreceptors (yR7) and yellow Dm8 neurons (yDm8) relies on the complementary expression of Dpr11 and DIP-γ, respectively, and the lack of either *dpr11* or *DIP-γ* leads to the failure of yR7 and yDm8 to recognize each other and subsequent yDm8 cell death (Carrillo et al., 2015; Courgeon and Desplan, 2019; Menon et al., 2019). Although extensive studies have uncovered roles for Dpr-DIP interactions, the regulation and downstream mechanisms are severely understudied.

Transcription factors (TFs) are the fate determinants of all cell types, and in neurons, they are master regulators of synaptic wiring by determining the expression of many factors including CSPs. In the fly olfactory system, cell type-specific expression of the TF *acj6* controls expression of a cell-surface code that instructs neurons to identify correct synaptic partners (Li et al., 2020; Xie et al., 2022). In the visual system, stochastic expression of Spineless (ss) determines the yR7 fate and controls the expression of *dpr11*, which is required for the connectivity with yellow Dm8s (Courgeon and Desplan, 2019). Similarly, during embryo development, several key TFs specify the neuroblast (NB) lineages, including *huckebein* (*hkb*), a TF expressed during embryo development (Brönner and Jäckle, 1991; Brönner and Jäckle, 1996; Gaul and Weigel, 1990; Weigel et al., 1990). In the developing nervous system, *hkb* is detected in 8 NB lineages and is required for the expression of the cell fate marker, *even-skipped* (*eve*), in NB4-2 (Broadus et al., 1995; Chu-LaGraff et al., 1995; McDonald and Doe, 1997). In *hkb* mutant embryos, RP2 MNs (also known as the dorsal Is MNs) derived from NB4-2 show severe wiring defects as they do not reach the correct muscles, suggesting that *hkb* controls specific CSPs for synaptic recognition in dorsal Is MNs (Chu-LaGraff et al., 1995).

In this study, we sought to identify genes involved in connectivity by developing a sensitized genetic background with known wiring CSPs that could be modified. Homozygous loss of *dpr10* and *DIP-α* led to complete loss of innervation of m4 by the dorsal Is MNs but *DIP-α*/+;*dpr10*/+ trans-heterozygous larvae showed a 50% reduction of m4-Is innervation frequency. This trans-heterozygous background was used for an F1 deficiency screen to identify dominant enhancers or suppressors of Dpr10/DIP-α-mediated connectivity. We screened deficiency lines from the Bloomington Deficiency Kit covering the right arm of the third chromosome (Cook et al., 2012; Roote and Russell, 2012), and within one interacting line, we identified *hkb* as a genetic regulator of *DIP-α*. *DIP-α* is expressed in both dorsal and ventral Is MNs, but interestingly, we found that *hkb* is only necessary for *DIP-α* expression and MN-muscle recognition in the dorsal Is MN, suggesting distinct regulatory mechanisms for *DIP-α* in dorsal and ventral Is MNs. Next, we showed that *hkb* functions through the dorsal Is MN specific TF, *eve*, as *DIP-α* expression and dorsal Is MN innervation are also disrupted in *eve* mutants. Genetic interaction tests between *hkb* and *eve* further confirmed that they act in the same pathway. In summary, our study suggests that Hkb acts through Eve and the downstream DIP-α to regulate MN-muscle connectivity and expands our understanding of how TFs instruct synaptic recognition.

## Results

### Genetic screen identifies *hkb*, a genetic interactor of Dpr10-DIP-α pathway

The *Drosophila* larval neuromuscular system provides an ideal model to study genetic programs that instruct synaptic recognition due to the ease of genetic manipulation and the stereotyped connectivity patterns. Each larval body wall hemisegment is innervated by one ventral and one dorsal Is MN that connect to the ventral or dorsal muscle groups, respectively, in a stereotyped manner (Figure 1A). In a previous study, we showed that among all MNs, *DIP-α* is expressed exclusively in these two Is MNs, and its interacting partner, Dpr10, is expressed in a subset of muscles (Ashley et al., 2019; Wang et al., 2022). The interaction between Dpr10 and DIP-α is required for the recognition between dorsal Is MNs and several dorsal muscles. Specifically, loss of either *dpr10* or *DIP-α* leads to complete loss of dorsal Is MN innervation on m4, suggesting that Dpr10-DIP-α interaction is absolutely required for m4-Is innervation. This easily scorable phenotype prompted us to ask what other genes are involved in this Dpr10-DIP-α-dependent synaptic recognition. Because loss of either CSP results in complete loss of m4-Is and single heterozygotes have either no or very mild phenotypes (see below), we created a sensitized genetic background in which one copy of *dpr10* and *DIP-α* was removed, m4- Is innervation frequency is reduced to ∼51% (Figure 1B), compared to a 90% m4-Is innervation frequency in wild type animals (Ashley et al., 2019). We chose a *dpr10* CRISPR (*dpr10^CR^*) allele and a GAL4 insertion allele of *DIP-α* (*DIP-α-GAL4*) derived from a MiMIC line, which disrupts endogenous *DIP-α* transcription and translation. Together with an *UAS-2xEGFP* construct, this *DIP-α-GAL4* allele aids identification of Is MN axons and NMJs on different muscles since *DIP-α* is exclusively expressed in Is MNs (Figure 1B). The reduced m4-Is innervation frequency in the sensitized background allowed us to screen for genetic interactors of the Dpr10-DIP-α pathway by introducing other mutations – if a mutation exacerbates or suppresses the decreased Is MN innervation on m4, we hypothesize that the gene may be part of the Dpr10/DIP-α pathway.

**Figure 1.**
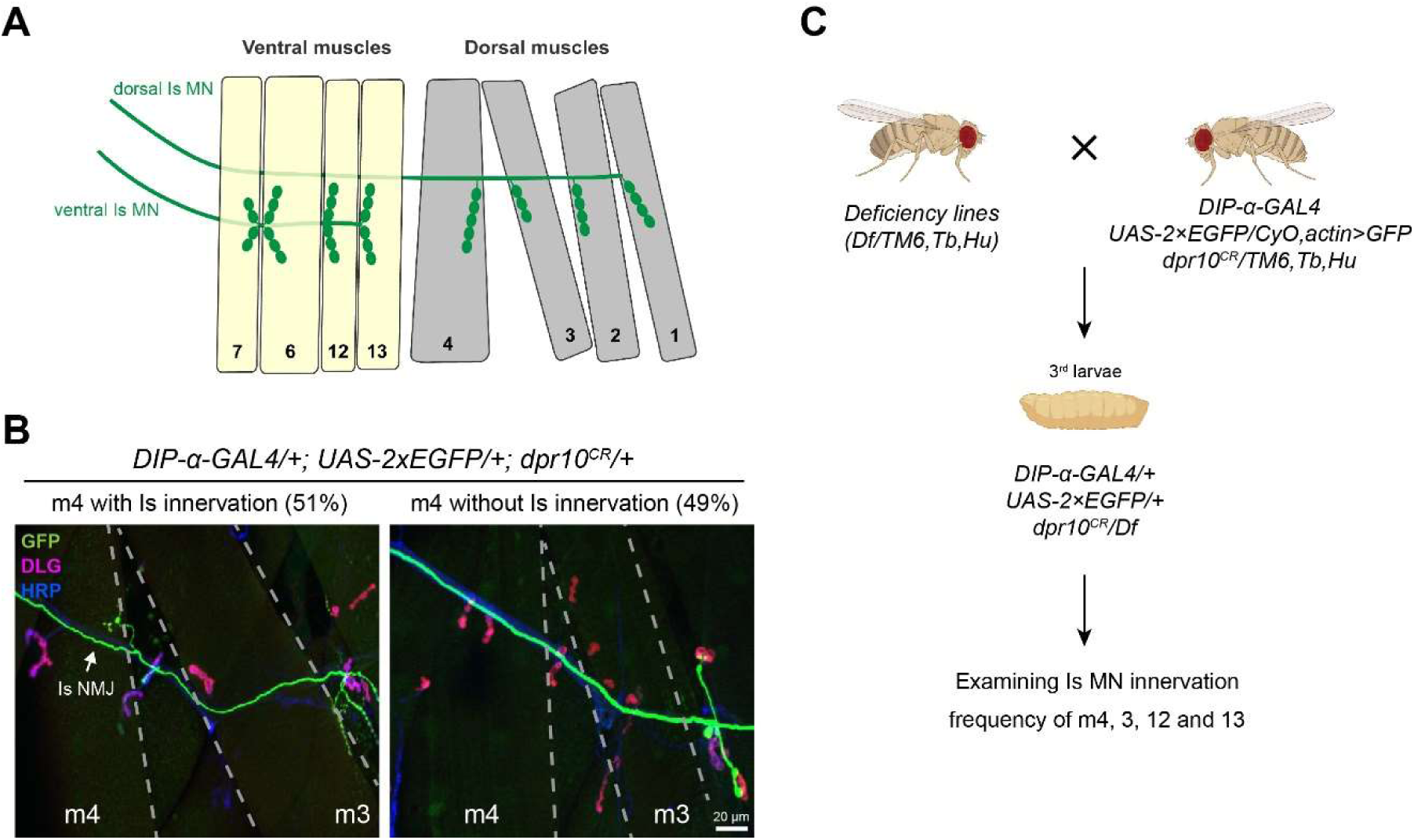
Establishing a sensitized background for deficiency screen. A. Cartoon depicting the innervation pattern of dorsal Is MN and ventral Is MN. B. Representative images of muscle 4 with Is innervation or without Is innervation, in trans-heterozygotes of *DIP-α* (the *DIP-α-GAL4* is also a null allele) and *dpr10*. 51% of m4 are innervated by the dorsal Is MN. GFP (green), DLG (magenta) and HRP (blue) are shown in the images. Arrow pointing to Is NMJ. C. Workflow of deficiency screen. Male flies carrying deficiency chromosome were crossed to females with *DIP-α* and *dpr10* mutations. Female third instar larvae were selected against the second and third chromosome balancers (*CyO,actin>GFP* and *TM6,Tb,Hu*). Triple-heterozygous larvae were dissected and Is innervation frequency on m4, 3, 12 and 13 were scored.

To improve the throughput, we utilized the Bloomington Deficiency (Df) kit and conducted an F1 dominant modifier screen (Figure 1C). We screened 105 Df lines that cover the entire *Drosophila* chromosome 3R. Each Df line was combined into the sensitized background to create triple heterozygotes and the m4-Is innervation frequency was quantified (Figure 2A). In addition, we also quantified the innervation frequency of Is MNs on other muscles, including dorsal muscle 3 (m3) and ventral muscle 12 (m12) and muscle 13 (m13) (Figure 2B-D). Although the Is innervation frequencies on these muscles were not significantly decreased in the sensitized background, we hypothesized that genetic interactors or redundant molecules of the Dpr10-DIP-α pathway may be uncovered. Compared to the sensitized background (red columns), we identified several Df lines that significantly increased or decreased Is innervation frequency (yellow columns) (Figure 2). The ED5100 Df line reduced m4-Is innervation frequency most significantly (p<0.01, Chi-square test), but did not affect Is innervation frequency on m12, m13 and m3 (p>0.05, Chi-square test), suggesting that it covers a gene(s) that may positively regulate the Dpr10-DIP-α pathway in the dorsal Is MNs for m4-Is recognition.

**Figure 2.**
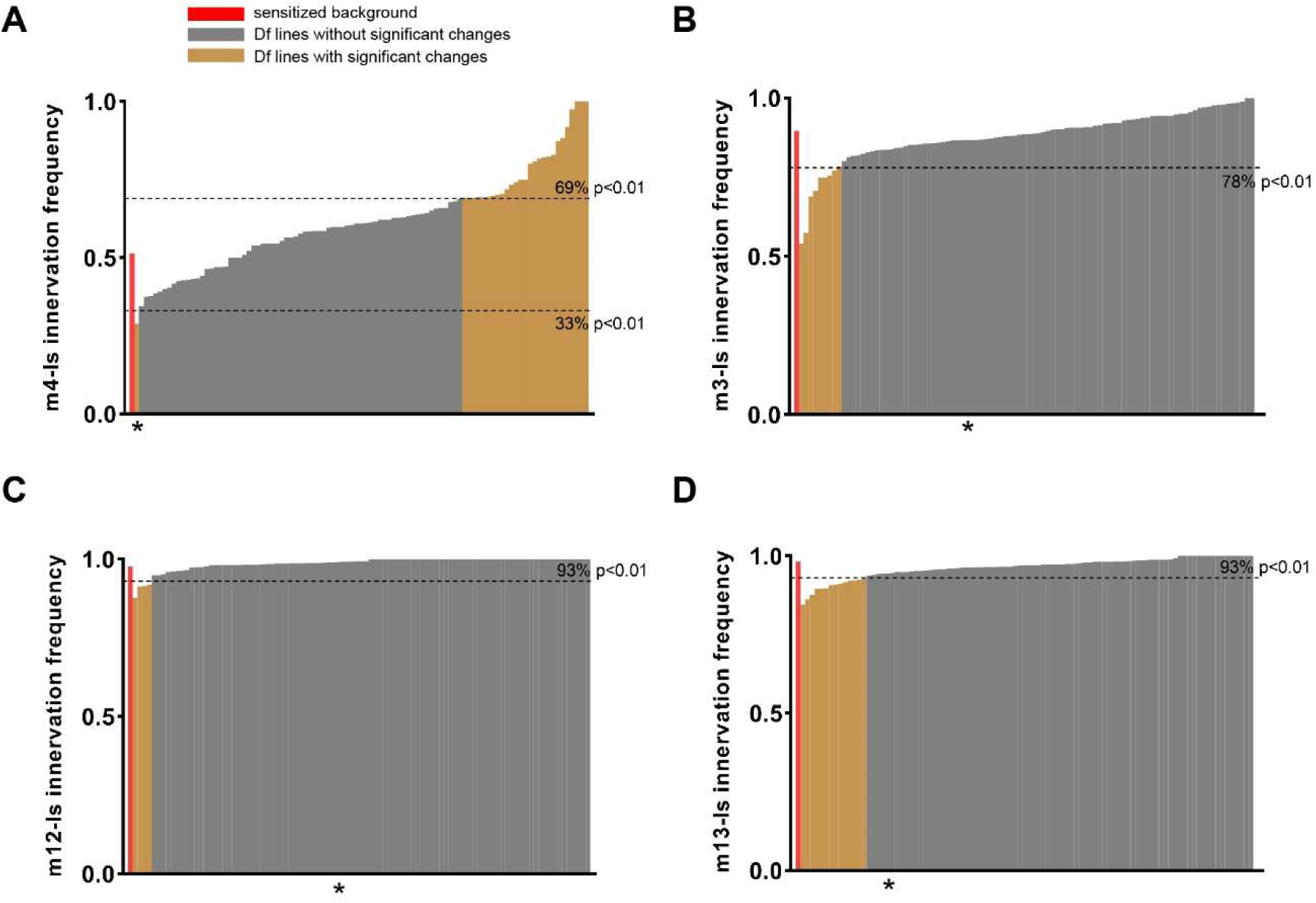
Deficiency screen revealed candidate regions that cover genetic interactors of *DIP-α* or *dpr10*. A-D. Is innervation frequency on (A) m4, (B) m3, (C) m12 and (D) m13. The red column indicates the control innervation frequency from the sensitized background (trans-heterozygotes of *DIP-α* and *dpr10*). Grey columns are non-significant from control whereas the yellow columns are the deficiency lines that show significantly different innervation frequencies compared to control. The cut-off p-values are indicated by dashed lines. Asterisk indicates ED5100.

ED5100 is a 900kb deletion that spans several genes and long non-coding RNAs. To narrow down the genomic region that covers our gene(s) of interest, we conducted a sub-screen using additional Df lines, ED5142 and ED5046, which partially overlap with the deletion in ED5100 (Figure 3A). We observed a similar decrease of m4- Is innervation frequency when the sensitized line was crossed to ED5046, but not to ED5142 (Figure 3B), suggesting that our gene(s) of interest is located in the region of ED5046 that overlaps with ED5100 but not ED5142, from 4,197 kb to 4,453 kb (Figure 3A). Several genes with known or potential neuronal functions are within this region, such as *auxilin*, *abstrakt*, *complexin*, *vps24*, *hkb*, *contactin*, *tube* and *lost*. We screened each candidate by combining a heterozygous mutant allele into our sensitized background and examined m4-Is innervation frequency. Of these candidates, we found that heterozygous loss of *hkb* (*hkb^2^/+*) exacerbated the m4-Is innervation defect when combined with the sensitized background (Figure 3C), suggesting that *hkb* genetically interacts with the Dpr10-DIP-α pathway.

**Figure 3.**
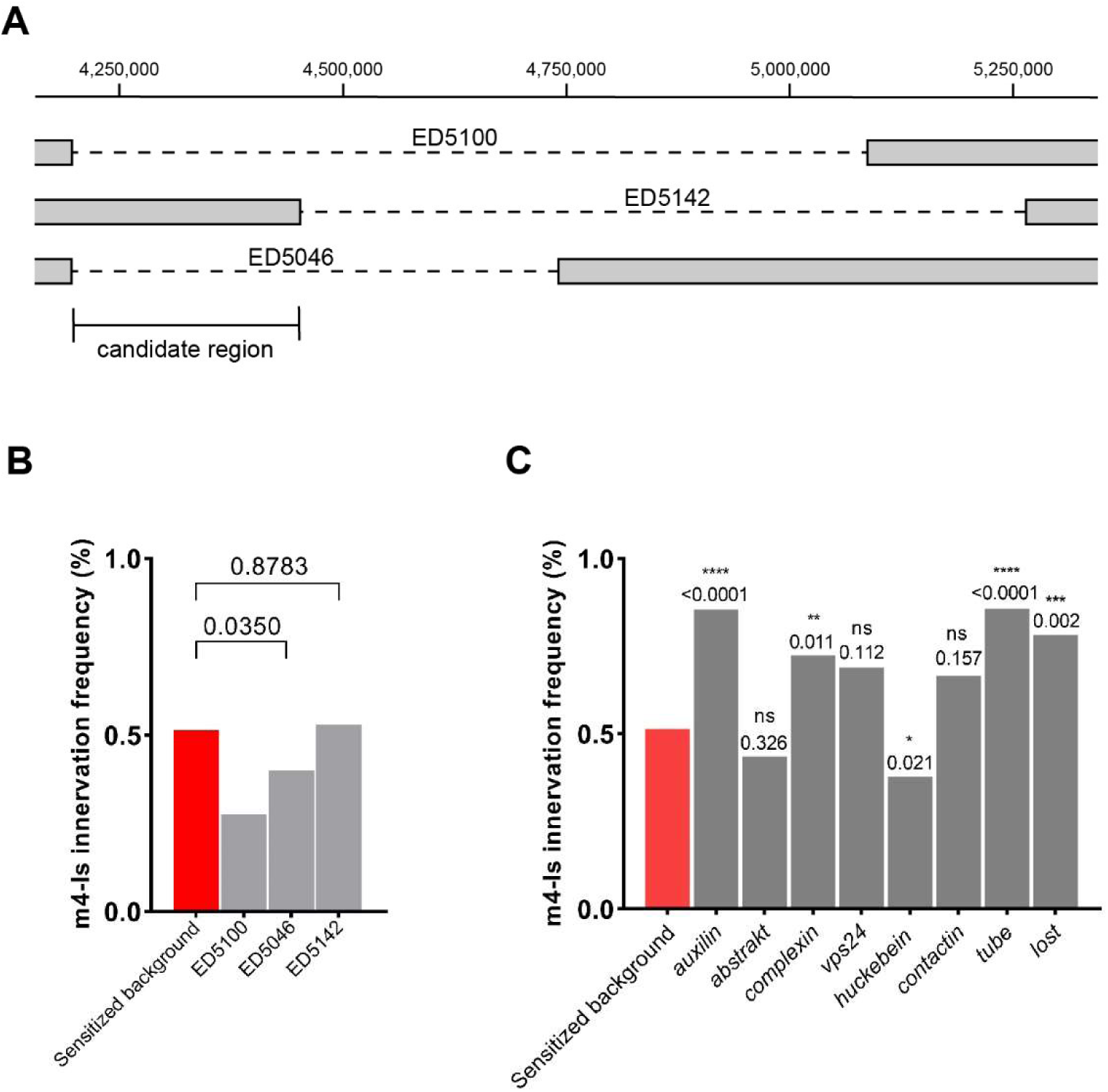
A sub-screen identified *huckebein* (*hkb)* as a genetic interactor. A. Cartoon depicting the deleted regions within deficiency lines ED5100, ED5142, and ED5046. B. Quantification shows a significant reduction of m4-Is innervation when combining ED5046, but not ED5142, with the sensitized background, suggesting the shared region between the original deficiency line (ED5100) and ED5046 covers the candidate gene(s). N (NMJs) = 177, 196, 155 and 155. p-values are indicted. C. Quantification of sub-screen of individual genes from candidate region shown in (A). Alleles used to create triple-heterozygotes are, *aux^D128^*, *abs^00620^*, *cpx^MI00784^*, *vps24^EY04708^*, *hkb^2^*, *cont^G5080^*, *tub^2^*, *lost^EY11645^*. Note that *huckebein* (*hkb^2^*) further reduced m4-Is innervation frequency. N (NMJs) = 177, 138, 62, 58, 29, 135, 39, 77, 78 and 57. p-values are indicted.

### *hkb* genetically interacts with *DIP-α*, but not *dpr10*

Our sensitized background is heterozygous for both *dpr10* and *DIP-α*. Therefore, *hkb* may genetically interact with either or both CSPs. Here, we examined genetic interaction between *hkb* and *dpr10* or *DIP-α* in trans-heterozygous animals. We combined two different *hkb* mutant alleles (*hkb^2^* or *hkb^A321R1^*) with a heterozygous *dpr10* mutant or *DIP-α* mutant and examined the m4-Is innervation frequency. In this and the following experiments, we used a *DIP-α* CRISPR (*DIP-α^CR^*) allele since we will primarily focus on m4-Is innervation and no longer need to identify Is NMJs on different muscles using *DIP-α-GAL4*.

In wild type animals, m4s are innervated about 90% of the time by the dorsal Is MN, and single heterozygotes of *dpr10* or *hkb* did not significantly decrease this innervation frequency (Figure 4A). We then examined trans-heterozygotes of *dpr10* and *hkb* and found that the m4-Is innervation frequency was not significantly changed compared to single heterozygotes (Figure 4A), suggesting that *hkb* is not a genetic interactor for *dpr10*. In contrast, heterozygous loss of *DIP-α* reduced m4-Is innervation frequency to 71% (Figure 4B), but the trans-heterozygotes of *DIP-α* and *hkb* further reduced the m4-Is innervation frequency to about 50%. Comparing the trans-heterozygous data with single heterozygotes (Figure 4B) suggests that *hkb* and *DIP-α* are in the same genetic pathway. Overall, these data suggest that *hkb* genetically interacts with *DIP-α* but not *dpr10*.

**Figure 4.**
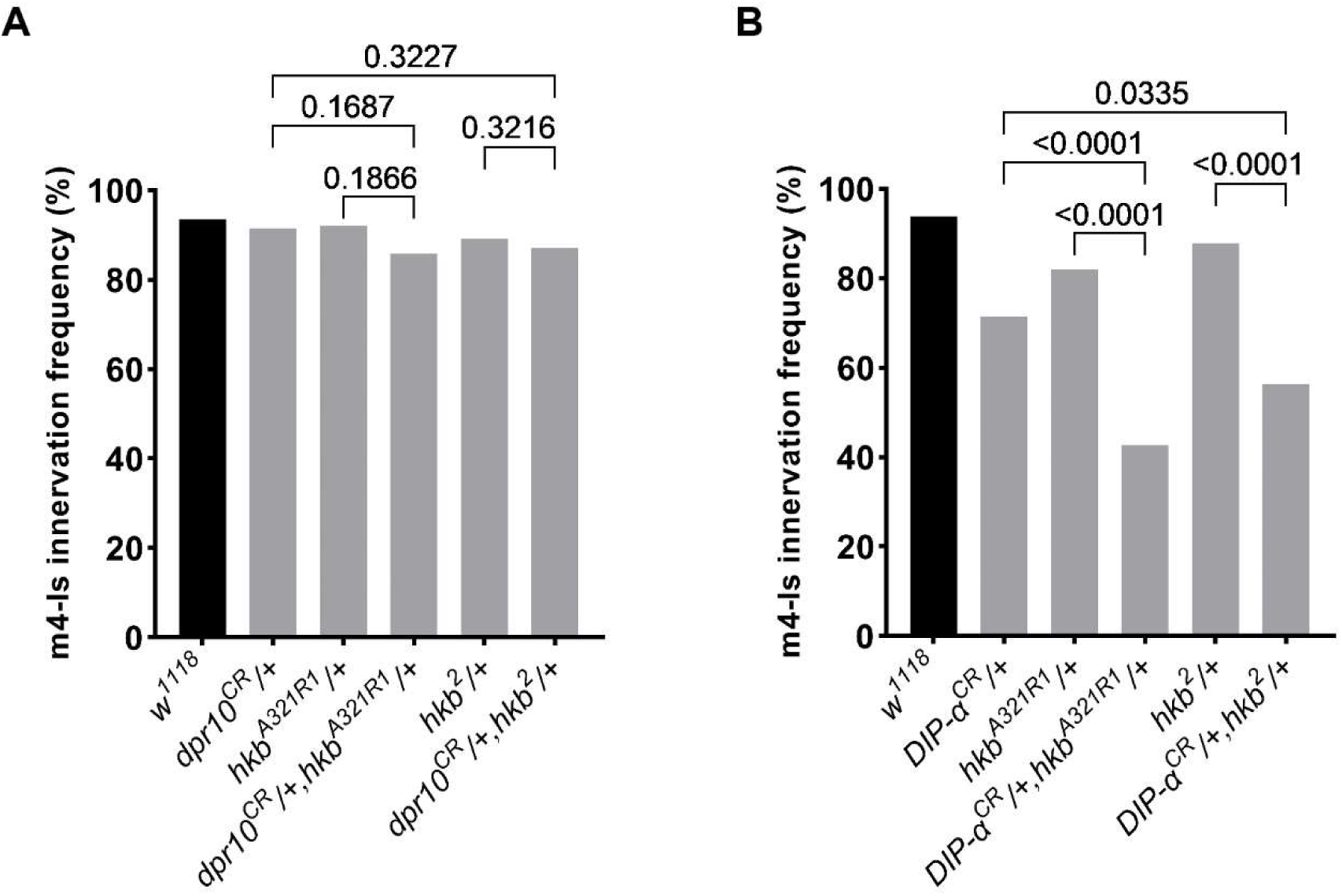
*hkb* genetically interacts with *DIP-α*, but not *dpr10*. A. Genetic interaction assay between *hkb* and *dpr10*. Single heterozygotes of *dpr10* or *hkb* did not have altered m4-Is innervation, and neither did the trans-heterozygotes. N (NMJs) = 93, 154, 83, 125, 102 and 149. p-values are indicted. B. Genetic interaction assay between *hkb* and *DIP-α*. Single heterozygotes of *DIP-α* or *hkb* had slightly decreased m4-Is innervation frequency, while the trans-heterozygotes showed a further reduction, suggesting that *hkb* genetically interacts with *DIP-α*. N (NMJs) = 114, 105, 106, 103, 99 and 103. p-values are indicted.

### *hkb* controls *DIP-α* expression in the dorsal Is MNs

Next, we asked how *hkb* genetically interacts with *DIP-α*. *DIP-α* is expressed in both dorsal and ventral Is MNs but not in muscles, and interestingly, prior studies found that *hkb* is expressed in NB4-2, which produces the dorsal Is MN (Chu-LaGraff et al., 1995; McDonald and Doe, 1997). Therefore, we wondered if the TF *hkb* is required for *DIP-α* expression in dorsal Is MNs. To visualize *DIP-α* expression, we used an endogenously tagged *DIP-α-EGFP* allele. In wild type animals, *DIP-α* is highly expressed in both dorsal and ventral Is MNs by stage 16 (Figure 5A, 5B and B’). However, in *hkb^2^* mutant embryos, expression of *DIP-α* in dorsal Is MNs is completely lost (Figure 5C). We confirmed loss of *DIP-α* in heteroallelic *hkb* mutant embryos (*hkb^2^*/*hkb^A321R1^*) (Figure 5D). Notably, the dorsal Is MN marker, *eve*, was also lost in *hkb* mutants as *hkb* is required for *eve* expression (Figure 5C and 5D) (Chu-LaGraff et al., 1995). These findings could be explained by the loss of the dorsal Is MNs; however, prior studies confirmed that dorsal Is MNs remain in *hkb* mutants even though they are *eve* negative (Chu-LaGraff et al., 1995; Fujioka et al., 2003). Therefore, the lack of *DIP-α-EGFP* is not due to missing Is MNs, but to the loss of *hkb*. In addition, as we hypothesized, *DIP-α-EGFP* is not affected in the ventral Is MNs in *hkb* mutants (Figure 5C’ and 5D’), suggesting that *hkb* only controls *DIP-α* expression in the dorsal Is MNs, which is consistent with the unchanged ventral Is MN innervation frequency for ED5100 in our genetic screen. Taken together, these data indicate that different mechanisms regulate the same CSP in different neurons, and *hkb* is required for *DIP-α* expression specifically in the dorsal Is MNs.

**Figure 5.**
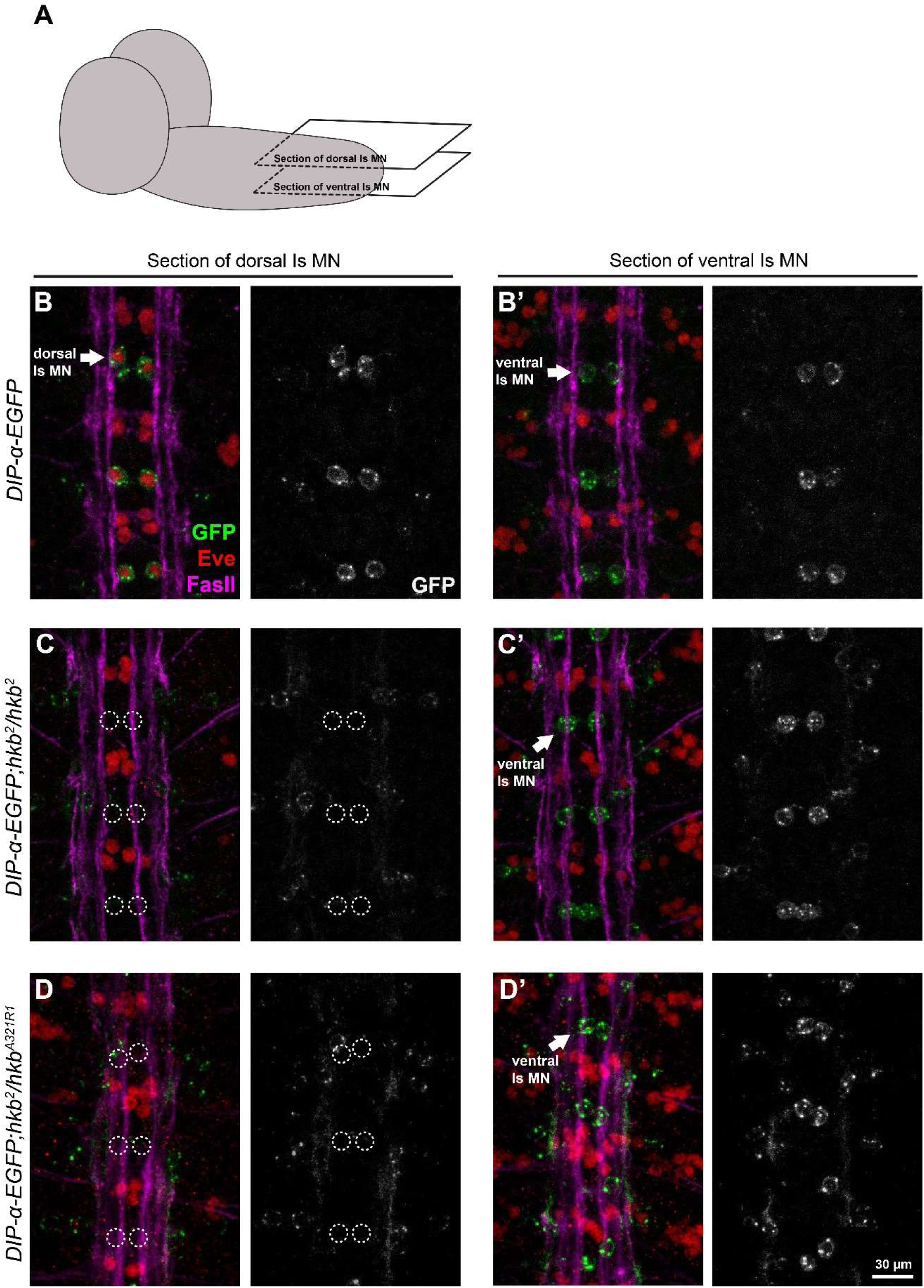
*hkb* is required for *DIP-α-EGFP* expression in dorsal Is MNs. A. Cartoon depicting the focal plane in B-D. B and B’. Representative images of dorsal Is MN cell bodies (arrows in B) and ventral Is MN cell bodies (arrows in B’) labeled with GFP (green), Eve (red) and FasII (magenta), in control embryos. *DIP-α-EGFP* is expressed in both dorsal and ventral Is MNs. C and C’. Representative images of dorsal Is MN cell bodies (dash circles in C) and ventral Is MN cell bodies (arrows in C’) in *hkb^2^* mutant embryos. Eve and DIP-α-EGFP are missing in dorsal Is MNs, whereas *DIP-α-EGFP* expression in ventral Is MNs is not affected. D and D’. Representative images of dorsal Is MN cell bodies (dash circles in D) and ventral Is MN cell bodies (arrows in D’) in heteroallelic *hkb* mutant embryos (*hkb^2^*/*hkb^A321R1^*). Eve and DIP-α-EGFP are missing in dorsal Is MNs, whereas *DIP-α-EGFP* expression in ventral Is MNs is not affected.

### *hkb* functions through *eve* to regulate *DIP-α* and dorsal Is MN innervation

As a TF, *hkb* may directly instruct *DIP-α* expression, or alternatively, function through other intermediate TFs. Prior studies reported that although *hkb* is expressed in the early NB4-2 that will give rise to dorsal Is MNs, its expression was turned off by stage 12, before synaptic recognition occurs in the neuromuscular system (Chu-LaGraff et al., 1995). However, a recent study found that *hkb* continues to be expressed in dorsal Is MNs during larval development (Jetti et al., 2023). Therefore, we decided to differentiate between the direct or indirect models of regulation. In NB4-2, a well-studied role of *hkb* is to trigger expression of the fate determinant TF, *eve*. Loss of *hkb* completely abolished *eve* expression in dorsal Is MNs (Figure 5) (Chu-LaGraff et al., 1995). Thus, we wondered if *hkb* functions through *eve* to regulate *DIP-α* expression. Utilizing the *DIP-α-EGFP* allele and a conditional *eve* knock-out which only lacks *eve* in dorsal Is MNs and siblings (Fujioka et al., 2003), we observed that *DIP-α* expression was lost in the dorsal Is MN (Figure 6A,B), and the expression in ventral Is MNs was not affected (Figure 6A’,B’). To further investigate the role of *eve* in synaptic recognition, we created *eve* and *DIP-α* trans-heterozygotes and examined m4-Is innervation frequency. Compared to single heterozygotes of *eve* or *DIP-α*, trans-heterozygotes significantly reduced m4-Is innervation frequency to about 60%, confirming that *eve* regulates m4-Is innervation by driving *DIP-α* expression in the dorsal Is MNs (Figure 6C). Finally, we examined the trans-heterozygotes of *hkb* and *eve* and found that partial loss of both *hkb* and *eve* reduced m4-Is innervation frequency to 75%, suggesting that *hkb* and *eve* indeed act in the same pathway to control synaptic recognition of the dorsal Is MNs (Figure 6D). Taken together, our results reveal a transcriptional cascade that regulates expression of wiring CSPs to guide MN-muscle recognition.

**Figure 6.**
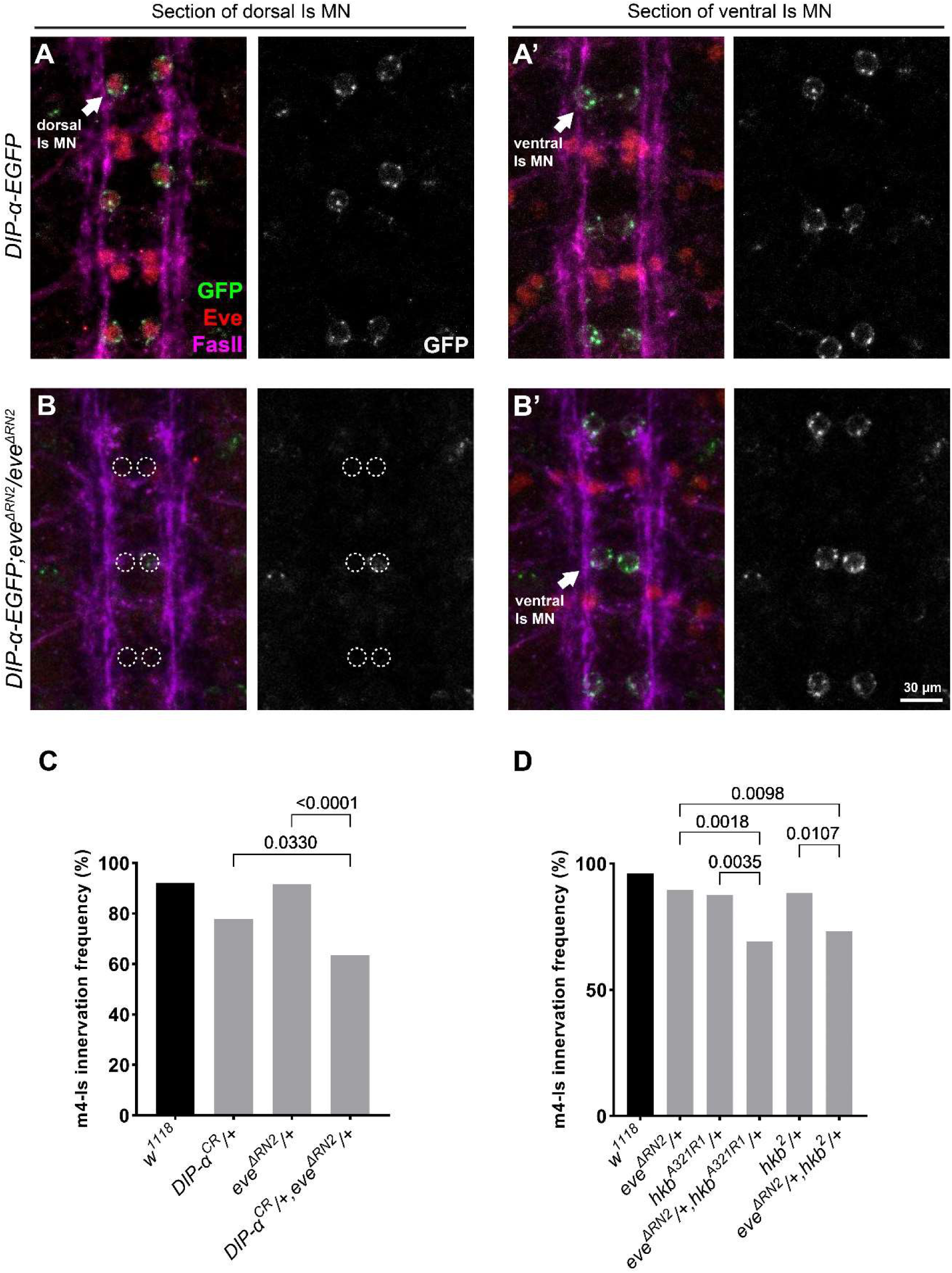
*hkb* functions through *eve* to regulate *DIP-α-EGFP* expression. A and A’. Representative images of dorsal Is MN cell bodies (arrows in A) and ventral Is MN cell bodies (arrows in A’) labeled with GFP (green), Eve (red) and FasII (magenta), in control embryos. *DIP-α-EGFP* is expressed in both dorsal and ventral Is MNs. B and B’. Representative images of dorsal Is MN cell bodies (dash circles in B) and ventral Is MN cell bodies (arrows in B’) in *eve^ΔRN2^* mutant embryos. Eve and DIP-α-EGFP are missing in dorsal Is MNs, whereas *DIP-α-EGFP* expression in ventral Is MNs is not affected. C. Genetic interaction assay between *eve* and *DIP-α*. Single heterozygotes of *eve* or *DIP-α* did not have altered m4-Is innervation, while the trans-heterozygotes showed a further reduction, suggesting that *eve* genetically interacts with *DIP-α*. N (NMJs) = 102, 104, 84 and 104. p-values are indicted. D. Genetic interaction assay between *hkb* and *eve*. Single heterozygotes of *hkb* or *eve* did not have altered m4-Is innervation, while the trans-heterozygotes showed a further reduction, suggesting that *hkb* genetically interacts with *eve* to regulate Is innervation. N (NMJs) = 103, 87, 105, 81, 112 and 86. p-values are indicted.

## Discussion

Synaptic recognition requires interactions of CSPs, and instrumental for proper circuit wiring is the expression of CSPs in specific cells. Underlying these precise expression patterns are transcriptional codes. Most studies have focused on the roles of TFs and CSPs independently, but less is known about the CSPs downstream of specific TFs in synaptic recognition. For example, *hkb* was implicated in pathfinding and target recognition of the dorsal Is MNs over three decades ago (Chu-LaGraff et al., 1995), but the molecular executors downstream of *hkb* were unknown. On the other hand, the well-studied Dpr10-DIP-α interaction was found to guide the recognition between the dorsal Is MN and m4, but the regulatory mechanisms controlling the expression of these CSPs were not known (Ashley et al., 2019). To identify additional components in the Dpr10-DIP-α pathway, including transcriptional regulators, we conducted a dominant modifier genetic screen and found that *hkb* is required for *DIP-α* expression in the dorsal Is MNs. Further examination revealed that *hkb* functions through *eve* to regulate *DIP-α* expression and dorsal Is MN innervation of m4, revealing a pathway linking TFs to specific CSPs and circuit assembly.

Interestingly, *hkb* is a gap gene originally implicated in embryo development. In the early embryo, three pattern organizing centers, the anterior, the posterior and the terminal, establish the anterior-posterior body plan by spatiotemporally regulating the expression of gap genes (Nasiadka et al., 2002). In the terminal control center, *torso* controls the expression of terminal gap genes including *hkb* (Weigel et al., 1990). *hkb* is expressed in the terminal cap during embryonic stages 5-6, and functions as a negative regulator to suppress gene expression in the terminal band, such as odd-paired (opa), Dichaete (D), caudal (cad) (Clark et al., 2022). Interestingly, in later embryonic stages, Hkb is expressed in a subset of neuroblasts where it is required for glial development (Iaco et al., 2006), serotoninergic neuron differentiation (Dittrich et al., 1997; Lundell et al., 1996), and for Eve expression to control motor axon pathfinding (Chu-LaGraff et al., 1995). However, the executors downstream of *hkb* for axon pathfinding are not known, and additionally, the role of *hkb* after the motor axon pathfinding stage has not been examined likely due to the lethality of null mutant embryos. In this study, we identified *hkb* as a *DIP-α* genetic interactor, and utilizing trans-heterozygous *hkb* hypomorph animals, we found that *hkb* is required for *DIP-α* expression to instruct innervation of the dorsal Is MN. Notably, our data suggests that *hkb* indirectly regulates *DIP-α* expression through the cell fate determinant TF, Eve. However, in a previous ChIP-Seq study profiling Eve target genes, many CSPs required for synaptic development were found, but *DIP-α* was not identified (Kudron et al., 2017). This could be due to *DIP-α* only expressed in a small subset of Eve-expressing cells, or more likely, because Eve also indirectly regulates *DIP-α* expression since it mostly functions as a transcriptional suppressor (Fujioka et al., 2003). Nevertheless, our genetic analyses focused on a single cell type and revealed regulatory relationships that would be obscured in sequencing-based profiling.

Excitatory MNs in *Drosophila* larvae are classified into type-Ib and type-Is MNs due to their terminal bouton size and innervation patterns. Notably, *DIP-α* is selectively expressed in the dorsal and ventral Is MNs but is absent in Ib MNs. We therefore initially hypothesized that a common regulatory program may be responsible for the expression of *DIP-α* in both Is MNs but absent in Ib MNs. However, we found that *hkb* and *eve* regulate *DIP-α* expression specifically in dorsal Is MNs. *hkb* and *eve* are not expressed in ventral Is MNs (derived from NB3-1) (McDonald and Doe, 1997), indicating distinct mechanisms regulate *DIP-α* expression in different MNs. Single-cell transcriptomics or candidate approaches in ventral Is MNs will aid in identifying other TFs that instruct *DIP-α* expression.

DIP-α is a member of the Dpr/DIP subfamilies of immunoglobulin CSPs. In a previous study, we mapped the expression of *dpr* and *DIP* genes in larval MNs and found that *dprs* were shared among many MNs and *DIPs* were more selectively expressed (Wang et al., 2022). Interestingly, each of the 33 MNs expresses a unique subset of *dprs* and *DIPs* to reveal a cell-specific cell surface code. These data suggest a highly complex regulatory transcriptional network is required to instruct the expression of these CSPs. An alternative but not mutually exclusive model is that one TF may regulate different CSPs in distinct neurons. A recent study in the fly olfactory circuit described a divergent “transcription factor to CSPs” relationship where the same TF, *acj6*, regulates many different CSPs in different cell types (Xie et al., 2022). Further identification of the TF network in the larval nervous system will help to understand how the TF code is transmitted into a CSP code to guide synaptic recognition.

In summary, the regulatory programs controlling expression of circuit wiring molecules are more complex than we hypothesized. The plethora of recent single-cell RNA-seq data will undoubtedly shed light on candidate TFs, but follow-up genetic analyses will be required to confirm causal relationships between TFs and CSPs that underlie circuit wiring.

## Methods

### Genetics

The following *Drosophila* lines were used in this study: *w^1118^* (Carrillo et al., 2015); *DIP-α-GAL4* (Ashley et al., 2019); *UAS-2×EGFP*; *dpr10^CR^*(Xu et al., 2018); *DIP-α^CR^* (Xu et al., 2018); *DIP-α-EGFP* (Tan et al., 2015); *hkb^A321R1^* (BL#2059) (Gaul and Weigel, 1990); *hkb^2^* (BL#5457) (Bossing et al., 1996); *eve^ΔRN2^*(Fujioka et al., 2003). Df lines discussed in this paper are: ED5100 (BL#9226), ED5046 (BL#9197), ED5142 (BL#9198). All lines used for screen and sub-screen are listed in Table 1 (Cook et al., 2012; Roote and Russell, 2012). For the genetic screen, males from the Df lines or mutant lines were crossed to sensitized females (*DIP-α-GAL4; UAS-2×EGFP/CyO,actin>GFP; dpr10^CR^/TM6,Tb,Hu*) and triple-heterozygous female larvae were selected to examine innervation frequency. For controls, *w^1118^* males were crossed to sensitized females to create trans-heterozygotes of *dpr10* and *DIP-α*.

**Table 1.**
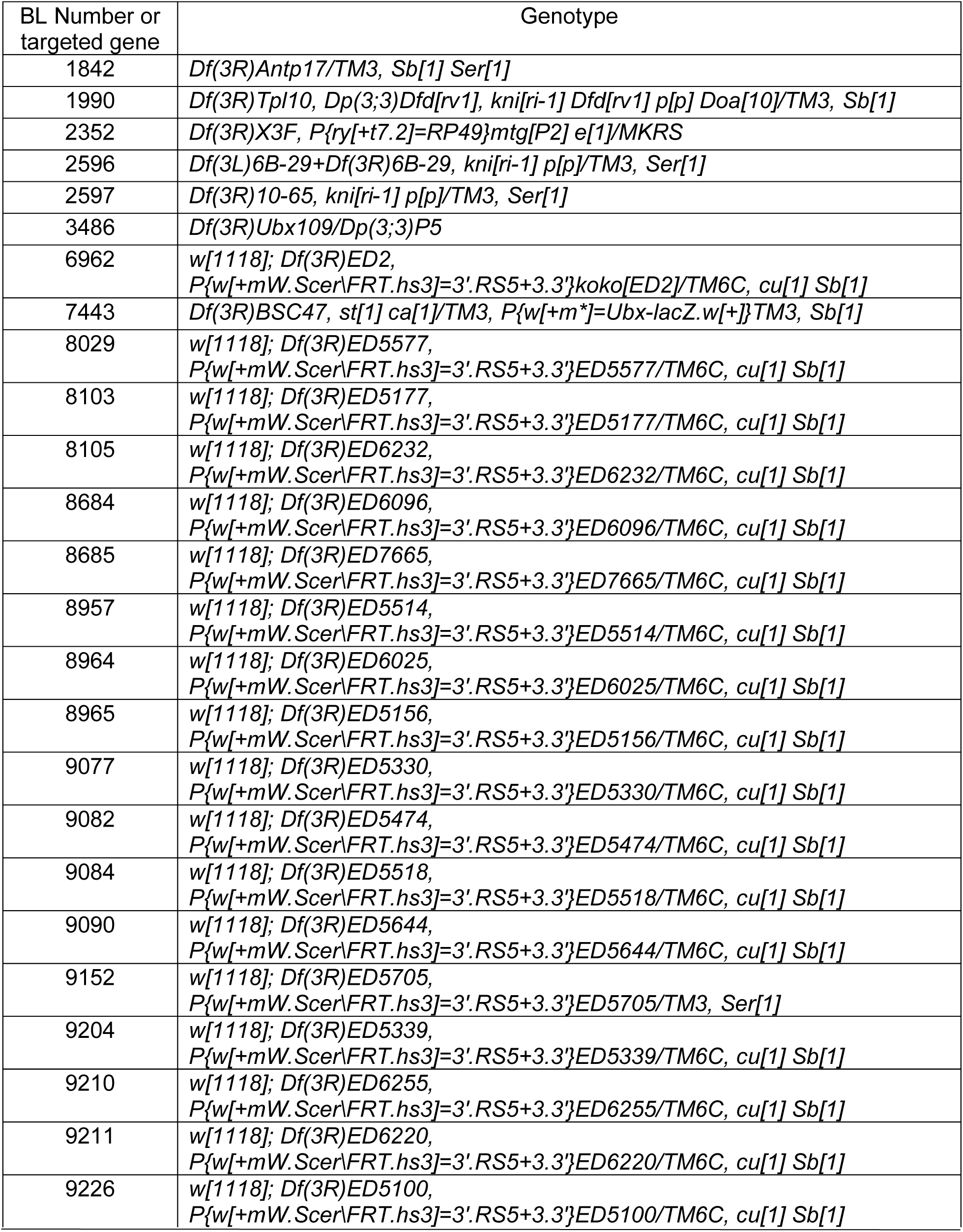
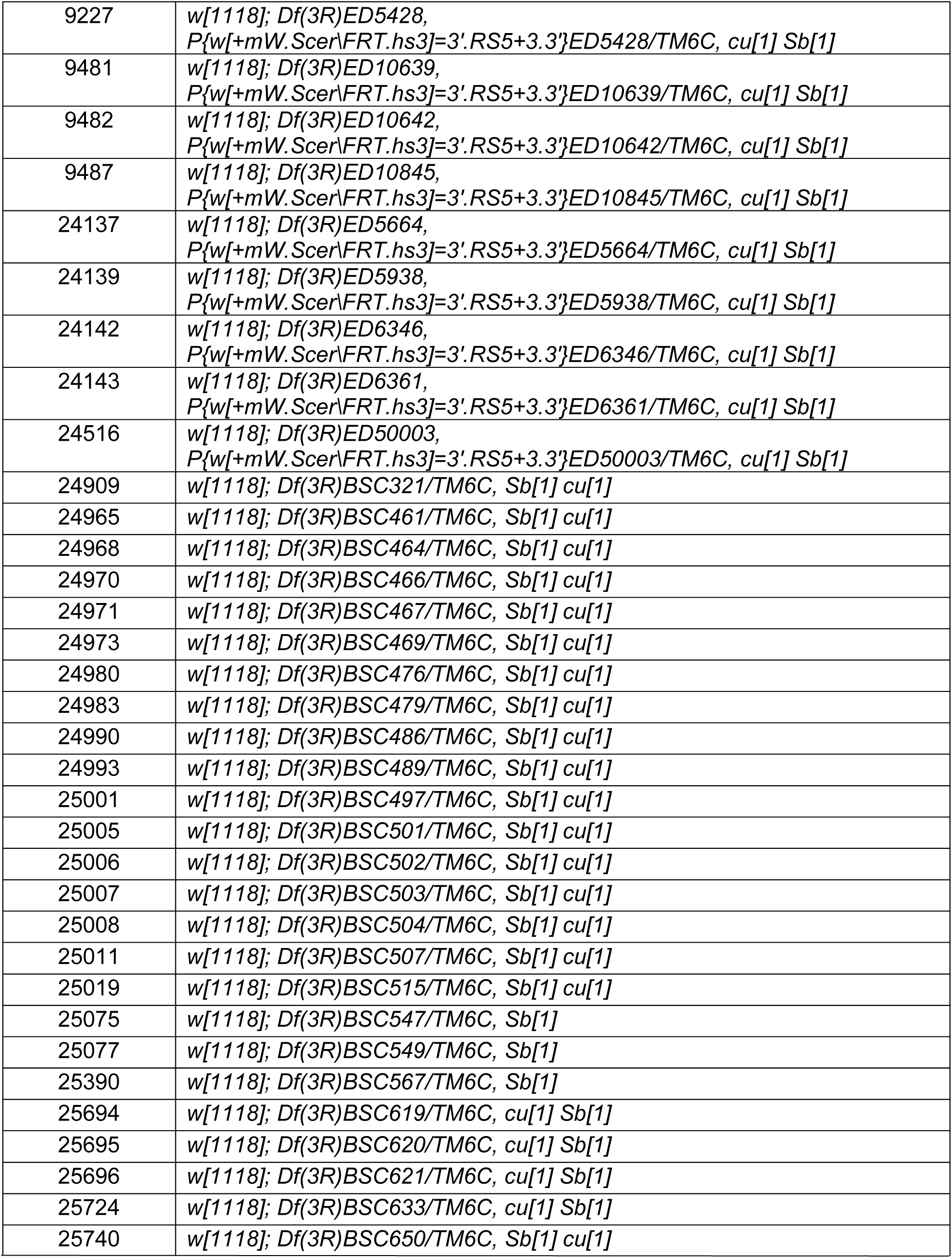
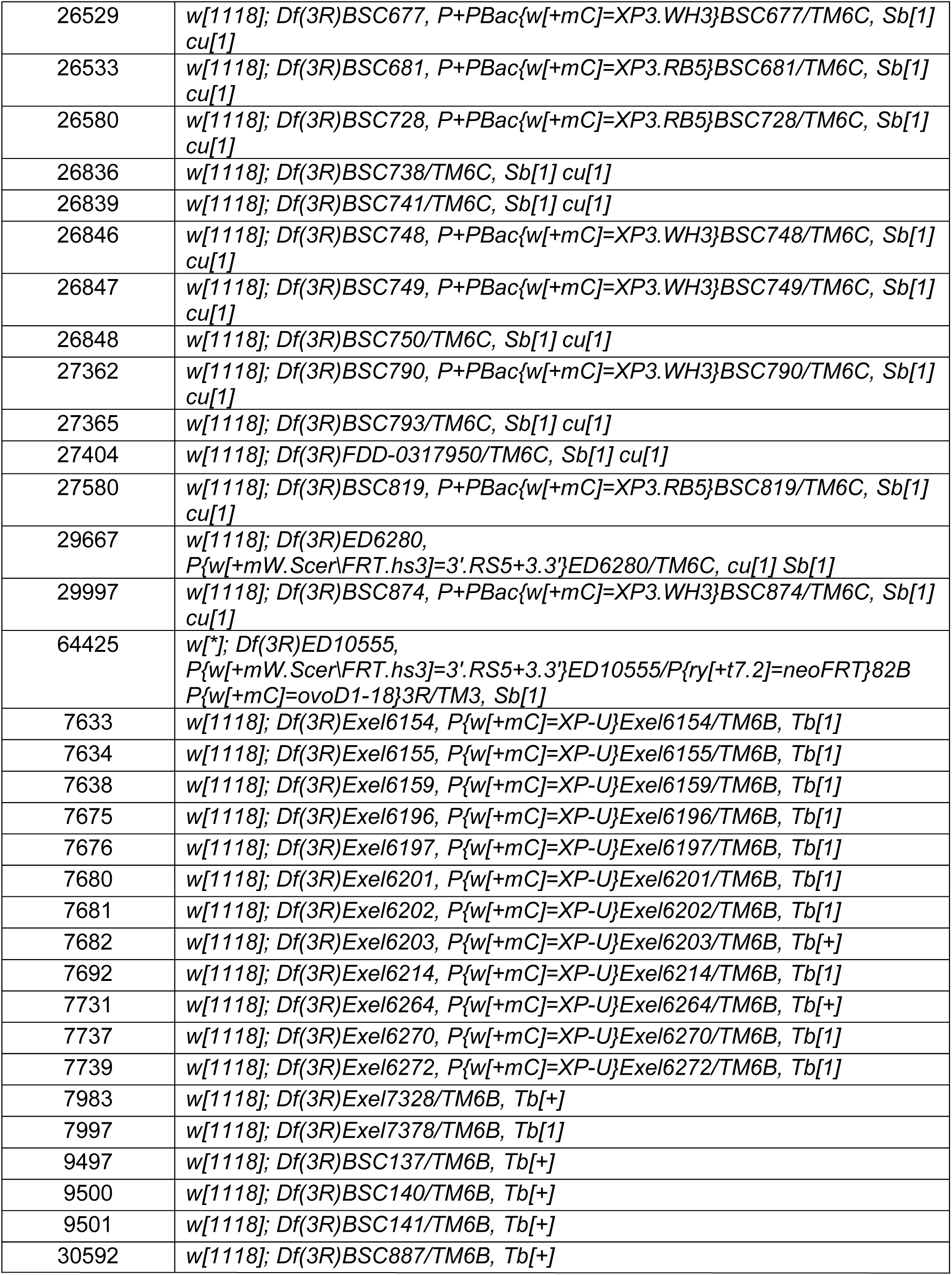
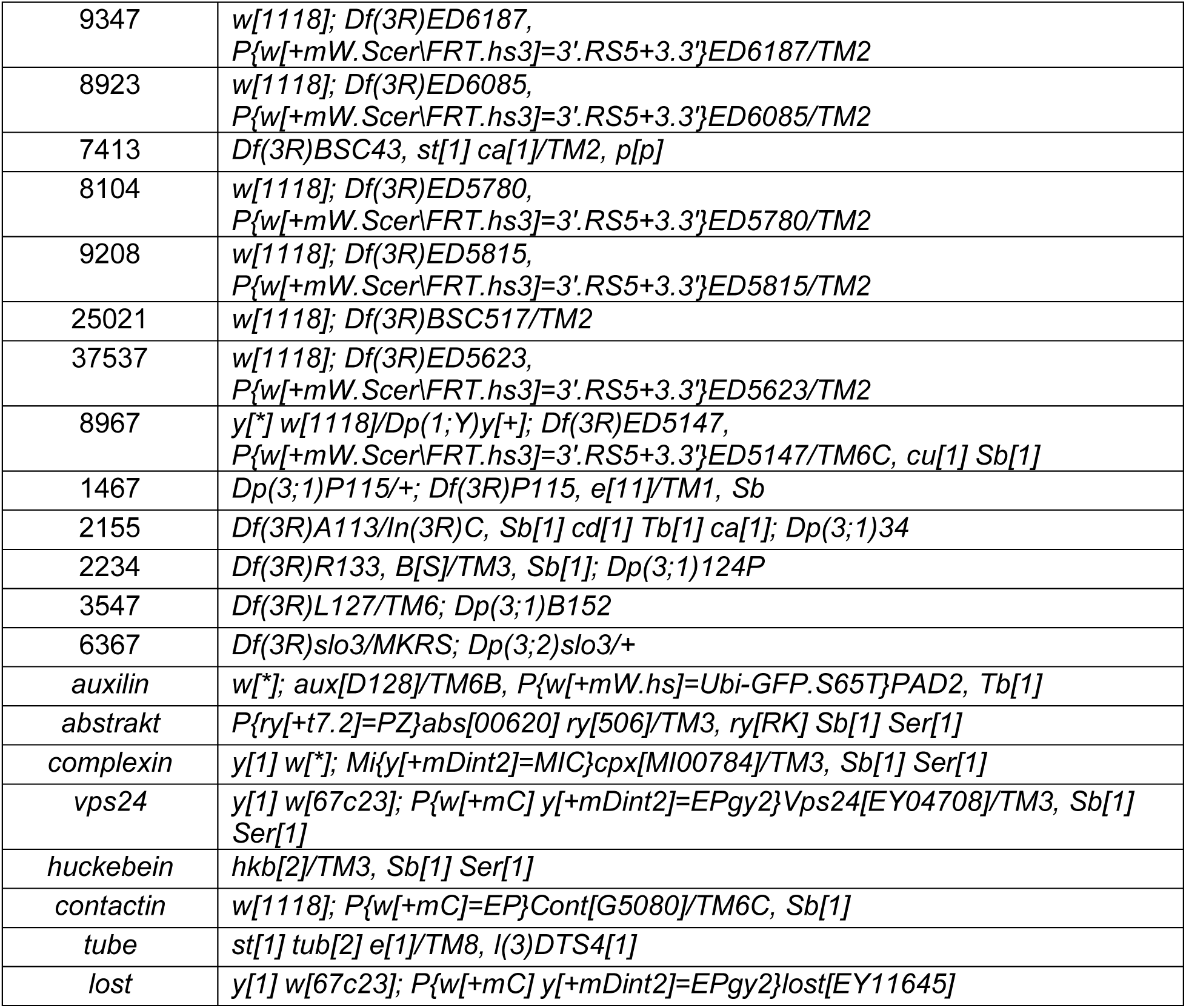
All Df lines and subscreen lines used in this study.

### Dissection, immunofluorescence and imaging

To examine Is MN innervation frequency, wandering third instar larvae were dissected as previously described (Wang et al., 2021). Briefly, larvae were collected and dissected on sylgard plates in PBS. Dissected fillets were fixed by 4% paraformaldehyde or Zamboni’s solution for 20 min at room temperature and then washed three times in PBT (PBS with 0.05% Triton X-100). Samples were then blocked for 1 h in 5% goat serum (5% goat serum diluted in PBT) and incubated with primary antibodies at 4 °C overnight. Primary antibodies were then removed, and samples were washed for three times before 2 h incubation at room temperature (or overnight at 4 °C) with secondary antibodies. Finally, secondary antibodies were washed out and samples were washed and mounted in Vectashield (Vector Laboratories).

To examine *DIP-α* expression, stage 16 embryos were dissected as previous described (Lobb-Rabe et al., 2022). Egg-laying chambers were set up with 40 females and 30 males and capped with grape juice plates. Eggs were collected for 2 h and grape juice plates covered in embryos were placed at 25 °C for 16 h for development. Embryos were staged under a Zeiss V20 stereoscope using the autofluorescence and morphology of the gut. Embryos at the correct stage were dechorionated and transferred onto a Superfrost Plus slides (Thermo Fisher Scientific, #22-037-246) covered by PBS. Embryos were then dissected with an electrolytically sharpened tungsten wire and stained similar to third instar samples. Antibodies used in this study were: rabbit anti-GFP (1:40k, gift from Michael Glozter, University of Chicago); rabbit anti-Eve (1:1000, gift from Ellie Heckscher, University of Chicago); mouse anti-DLG (1:100, Developmental Studies Hybridoma Bank [DSHB] #4F3); mouse anti-FasII (1:100, DSHB #1D4); goat anti-rabbit Alexa 488 (1:500, Invitrogen #A11008); goat anti-rabbit Alexa 568 (1:500, Invitrogen #A11036); goat anti-mouse Alexa 568 (1:500, Invitrogen #A11031); goat anti-mouse Alexa 647 (1:500, Invitrogen #A32728); goat anti-HRP Alexa 405 (1:100, Jackson Immunological Research #123-475-021); goat anti-HRP Alexa 647 (1:100, Jackson Immunological Research #123-605-021).

Images were acquired on a Zeiss LSM800 confocal microscope using a 40X plan-neofluar 1.3 NA objective, or a 63X plan-apo 1.4 NA objective. The same imaging parameters were applied to samples from the same set of experiments. Images were then analyzed and processed in ImageJ.

### Quantification of Is MN innervation frequency

To examine Is MN innervation frequency, at least 6 third instar larvae were dissected and stained with anti-GFP (marker for Is MNs), anti-DLG and anti-HRP. Samples were visualized under a Zeiss AxioImager M2 scope with a Lumen light engine with a 20× Plan Apo 0.8 NA objective or an Olympus BX43 with an X-Cite 120LEDmini LED fluorescent illuminator (will add objective). Is and Ib NMJs can be distinguished by bouton size, DLG intensity and whether it was GFP positive. If there was at least one Is bouton present on the muscle, it is scored as “innervated”, otherwise it is scored as “not innervated”. For each animal, m12, 13, 4 and 3 from abdominal hemisegments A2-A6 were assayed, and we collected a sample size of 50 – 60 hemisegments for each muscle. Innervation frequency was calculated as the percentage of “innervated” muscles in all muscles examined.

### Statistical analysis

As we were comparing the innervation frequency between two groups, we performed Chi-square test followed by Yates’ correction using Prism 8 software. Innervation frequencies and p-values were reported in the figure legends.

## Acknowledgments

This work is supported by NSF IOS-2048080, NINDS R01 NS123439, and a UChicago Faculty Diversity Grant to R.A.C., by NINDS R15 NS101692 01 & 02 to C.G.V., and by NIH T32 GM139782 to R.Z. and L.S. This work is also supported by funds from the UChicago Biological Science Division, Committee of Developmental Biology and Department of Molecular Genetics & Cellular Biology, and by the Skidmore College Summer Collaborative Research Program. We thank the Bloomington Drosophila Stock Center (NIH P40OD018537) for fly lines. The monoclonal antibodies 4F3 and 1D4 were developed by Goodman, C. and they were obtained from the Developmental Studies Hybridoma Bank, created by the NICHD of the NIH and maintained at the University of Iowa, Department of Biology. We would like to thank Ellie Hecksher and Michael Glotzer for sharing resources. We thank the Skidmore Microscopy Imaging Center (SMIC) for the use of their microscopy resources. For their contributions to the screen, we also would like to thank the Spring 2018 Skidmore Neurophysiology students (Emily Blunt, Haoyang Huang, Jessy Idemoto, Dilhan Sirtalan, Julie Wang, Rob Warden, and Mary Beth Zahnleuter) and the Spring 2022 Skidmore Neurophysiology students (Jessica Auerbach, Margaret Besthoff, Gene Choi, Joa Comellas, DJ Flam, Melaina Gilbert, Kaylee Hua, Alexander Nardone, Bryan Taylor, and Victoria Thorpe). We would also like to thank Richard Fehon, David Pincus, and members from the Carrillo laboratory for valuable discussions and comments. Some cartoon representations were generated by BioRender.

## Author contribution

Y.W. and R.A.C designed research; Y.W., R.S., V.S., L.S., T.J.G., and B.W. performed experiments; Y.W., R.S. and J.A. analyzed data; Y.W. wrote the manuscript; J.A., C.G.V., and R.A.C. edited the manuscript.

## References

Ashley, J., Sorrentino, V., Lobb-Rabe, M., Nagarkar-Jaiswal, S., Tan, L., Xu, S., Xiao, Q., Zinn, K. and Carrillo, R. A. (2019). Transsynaptic interactions between IgSF proteins DIP-α and Dpr10 are required for motor neuron targeting specificity. eLife 8, e42690.

Barish, S., Nuss, S., Strunilin, I., Bao, S., Mukherjee, S., Jones, C. D. and Volkan, P. C. (2018). Combinations of DIPs and Dprs control organization of olfactory receptor neuron terminals in Drosophila. Plos Genet 14, e1007560.

Bornstein, B., Meltzer, H., Adler, R., Alyagor, I., Berkun, V., Cummings, G., Reh, F., Keren-Shaul, H., David, E., Riemensperger, T., et al. (2021). Transneuronal Dpr12/DIP-δ interactions facilitate compartmentalized dopaminergic innervation of Drosophila mushroom body axons. Embo J e105763.

Bossing, T., Technau, G. M. and Doe, C. Q. (1996). huckebein is required for glial development and axon pathfinding in the neuroblast 1-1 and neuroblast 2-2 lineages in the Drosophila central nervous system. Mech Develop 55, 53–64.

Broadus, J., Skeath, J. B., Spana, E. P., Bossing, T., Technau, G. and Doe, C. Q. (1995). New neuroblast markers and the origin of the aCC/pCC neurons in the Drosophila central nervous system. Mech Develop 53, 393–402.

Brönner, G. and Jäckle, H. (1991). Control and function of terminal gap gene activity in the posterior pole region of the Drosophila embryo. Mech Develop 35, 205–211.

Brönner, G. and Jäckle, H. (1996). Regulation and function of the terminal gap gene huckebein in the Drosophila blastoderm. Int J Dev Biology 40, 157–65.

Brovero, S. G., Fortier, J. C., Hu, H., Lovejoy, P. C., Newell, N. R., Palmateer, C. M., Tzeng, R.-Y., Lee, P.-T., Zinn, K. and Arbeitman, M. N. (2021). Investigation of Drosophila fruitless neurons that express Dpr/DIP cell adhesion molecules. Elife 10, e63101.

Carrillo, R. A., Özkan, E., Menon, K. P., Nagarkar-Jaiswal, S., Lee, P.-T., Jeon, M., Birnbaum, M. E., Bellen, H. J., Garcia, K. C. and Zinn, K. (2015). Control of Synaptic Connectivity by a Network of Drosophila IgSF Cell Surface Proteins. Cell 163, 1770–1782.

Cheng, S., Ashley, J., Kurleto, J. D., Lobb-Rabe, M., Park, Y. J., Carrillo, R. A. and Özkan, E. (2019). Molecular basis of synaptic specificity by immunoglobulin superfamily receptors in Drosophila. Elife 8, e41028.

Chiba, A., Snow, P., Keshishian, H. and Hotta, Y. (1995). Fasciclin III as a synaptic target recognition molecule in Drosophila. Nature 374, 166–168.

Chu-LaGraff, Q., Schmid, A., Leidel, J., Brönner, G., Jäckle, H. and Doe, C. Q. (1995). huckebein specifies aspects of CNS precursor identity required for motoneuron axon pathfinding. Neuron 15, 1041–1051.

Clark, E., Battistara, M. and Benton, M. A. (2022). A timer gene network is spatially regulated by the terminal system in the Drosophila embryo. Elife 11,.

Cook, R. K., Christensen, S. J., Deal, J. A., Coburn, R. A., Deal, M. E., Gresens, J. M., Kaufman, T. C. and Cook, K. R. (2012). The generation of chromosomal deletions to provide extensive coverage and subdivision of the Drosophila melanogaster genome. Genome Biology 13, R21.

Cosmanescu, F., Katsamba, P. S., Sergeeva, A. P., Ahlsen, G., Patel, S. D., Brewer, J. J., Tan, L., Xu, S., Xiao, Q., Nagarkar-Jaiswal, S., et al. (2018). Neuron-Subtype-Specific Expression, Interaction Affinities, and Specificity Determinants of DIP/Dpr Cell Recognition Proteins. Neuron 100, 1385–1400.e6.

Courgeon, M. and Desplan, C. (2019). Coordination between stochastic and deterministic specification in the Drosophila visual system. Science 366, eaay6727.

Davis, G. W., Schuster, C. M. and Goodman, C. S. (1997). Genetic Analysis of the Mechanisms Controlling Target Selection: Target-Derived Fasciclin II Regulates the Pattern of Synapse Formation. Neuron 19, 561–573.

Dittrich, R., Bossing, T., Gould, A. P., Technau, G. M. and Urban, J. (1997). The differentiation of the serotonergic neurons in the Drosophila ventral nerve cord depends on the combined function of the zinc finger proteins Eagle and Huckebein. Dev Camb Engl 124, 2515–25.

Fujioka, M., Lear, B. C., Landgraf, M., Yusibova, G. L., Zhou, J., Riley, K. M., Patel, N. H. and Jaynes, J. B. (2003). Even-skipped, acting as a repressor, regulates axonal projections in Drosophila. Development 130, 5385–5400.

Garrett, A. M., Khalil, A., Walton, D. O. and Burgess, R. W. (2018). DSCAM promotes self-avoidance in the developing mouse retina by masking the functions of cadherin superfamily members. Proc National Acad Sci 115, 201809430.

Gaul, U. and Weigel, D. (1990). Regulation of Krüppel expression in the anlage of the Malpighian tubules in the Drosophila embryo. Mech Develop 33, 57–67.

Hattori, D., Chen, Y., Matthews, B. J., Salwinski, L., Sabatti, C., Grueber, W. B. and Zipursky, S. L. (2009). Robust discrimination between self and non-self neurites requires thousands of Dscam1 isoforms. Nature 461, 644–648.

Hoang, B. and Chiba, A. (2001). Single-Cell Analysis of Drosophila Larval Neuromuscular Synapses. Dev Biol 229, 55–70.

Iaco, R. D., Soustelle, L., Kammerer, M., Sorrentino, S., Jacques, C. and Giangrande, A. (2006). Huckebein-mediated autoregulation of Glide/Gcm triggers glia specification. Embo J 25, 244– 254.

Jetti, S. K., Crane, A. B., Akbergenova, Y., Aponte-Santiago, N. A., Cunningham, K. L., Whittaker, C. A. and Littleton, J. T. (2023). Molecular Logic of Synaptic Diversity Between Drosophila Tonic and Phasic Motoneurons. bioRxiv 2023.01.17.524447.

Kose, H., Rose, D., Zhu, X. and Chiba, A. (1997). Homophilic synaptic target recognition mediated by immunoglobulin-like cell adhesion molecule Fasciclin III. Dev Camb Engl 124, 4143–52.

Krishnaswamy, A., Yamagata, M., Duan, X., Hong, Y. K. and Sanes, J. R. (2015). Sidekick 2 directs formation of a retinal circuit that detects differential motion. Nature 524, 466–470.

Kudron, M. M., Victorsen, A., Gevirtzman, L., Hillier, L. W., Fisher, W. W., Vafeados, D., Kirkey, M., Hammonds, A. S., Gersch, J., Ammouri, H., et al. (2017). The modERN Resource: Genome-Wide Binding Profiles for Hundreds of Drosophila and Caenorhabditis elegans Transcription Factors. Genetics 208, genetics.300657.2017.

Li, H., Li, T., Horns, F., Li, J., Xie, Q., Xu, C., Wu, B., Kebschull, J. M., McLaughlin, C. N., Kolluru, S. S., et al. (2020). Single-Cell Transcriptomes Reveal Diverse Regulatory Strategies for Olfactory Receptor Expression and Axon Targeting. Curr Biol 30, 1189–1198.e5.

Lnenicka, G. A. and Keshishian, H. (2000). Identified motor terminals in Drosophila larvae show distinct differences in morphology and physiology. J Neurobiol 43, 186–197.

Lobb-Rabe, M., DeLong, K., Salazar, R. J., Zhang, R., Wang, Y. and Carrillo, R. A. (2022). Dpr10 and Nocte are required for Drosophila motor axon pathfinding. Neural Dev 17, 10.

Lundell, M. J., Chu-LaGraff, Q., Doe, C. Q. and Hirsh, J. (1996). TheengrailedandhuckebeinGenes Are Essential for Development of Serotonin Neurons in theDrosophilaCNS. Mol Cell Neurosci 7, 46–61.

McDonald, J. A. and Doe, C. Q. (1997). Establishing neuroblast-specific gene expression in the Drosophila CNS: huckebein is activated by Wingless and Hedgehog and repressed by Engrailed and Gooseberry. Dev Camb Engl 124, 1079–87.

Menon, K. P., Kulkarni, V., Takemura, S., Anaya, M. and Zinn, K. (2019). Interactions between Dpr11 and DIP-γ control selection of amacrine neurons in Drosophila color vision circuits. Elife 8, e48935.

Nasiadka, A., Dietrich, B. H. and Krause, H. M. (2002). Anterior-posterior patterning in the Drosophila embryo. Adv. Dev. Biology Biochem. 12, 155–204.

Özkan, E., Carrillo, R. A., Eastman, C. L., Weiszmann, R., Waghray, D., Johnson, K. G., Zinn, K., Celniker, S. E. and Garcia, K. C. (2013). An Extracellular Interactome of Immunoglobulin and LRR Proteins Reveals Receptor-Ligand Networks. Cell 154, 228–239.

Roote, J. and Russell, S. (2012). Toward a complete Drosophiladeficiency kit. Genome Biology 13, 149.

Sergeeva, A. P., Katsamba, P. S., Cosmanescu, F., Brewer, J. J., Ahlsen, G., Mannepalli, S., Shapiro, L. and Honig, B. (2020). DIP/Dpr interactions and the evolutionary design of specificity in protein families. Nat Commun 11, 2125.

Shen, K. and Bargmann, C. I. (2003). The Immunoglobulin Superfamily Protein SYG-1 Determines the Location of Specific Synapses in C. elegans. Cell 112, 619–630.

Shen, K., Fetter, R. D. and Bargmann, C. I. (2004). Synaptic Specificity Is Generated by the Synaptic Guidepost Protein SYG-2 and Its Receptor, SYG-1. Cell 116, 869–881.

Tan, L., Zhang, K. X., Pecot, M. Y., Nagarkar-Jaiswal, S., Lee, P.-T., Takemura, S., McEwen, J. M., Nern, A., Xu, S., Tadros, W., et al. (2015). Ig Superfamily Ligand and Receptor Pairs Expressed in Synaptic Partners in Drosophila. Cell 163, 1756–69.

Venkatasubramanian, L., Guo, Z., Xu, S., Tan, L., Xiao, Q., Nagarkar-Jaiswal, S. and Mann, R. S. (2019). Stereotyped terminal axon branching of leg motor neurons mediated by IgSF proteins DIP-α and Dpr10. Elife 8, e42692.

Wang, J., Ma, X., Yang, J. S., Zheng, X., Zugates, C. T., Lee, C.-H. J. and Lee, T. (2004). Transmembrane/Juxtamembrane Domain-Dependent Dscam Distribution and Function during Mushroom Body Neuronal Morphogenesis. Neuron 43, 663–672.

Wang, Y., Lobb-Rabe, M., Ashley, J., Anand, V. and Carrillo, R. A. (2021). Structural and functional synaptic plasticity induced by convergent synapse loss in the Drosophila neuromuscular circuit. J Neurosci JN-RM-1492–20.

Wang, Y., Lobb-Rabe, M., Ashley, J., Chatterjee, P., Anand, V., Bellen, H. J., Kanca, O. and Carrillo, R. A. (2022). Systematic expression profiling of dprs and DIPs reveals cell surface codes in Drosophila larval motor and sensory neurons. Development 149,.

Weigel, D., Jürgens, G., Klingler, M. and Jäckle, H. (1990). Two Gap Genes Mediate Maternal Terminal Pattern Information in Drosophila. Science 248, 495–498.

Winberg, M. L., Mitchell, K. J. and Goodman, C. S. (1998). Genetic Analysis of the Mechanisms Controlling Target Selection: Complementary and Combinatorial Functions of Netrins, Semaphorins, and IgCAMs. Cell 93, 581–591.

Xie, Q., Li, J., Li, H., Udeshi, N. D., Svinkina, T., Orlin, D., Kohani, S., Guajardo, R., Mani, D. R., Xu, C., et al. (2022). Transcription factor Acj6 controls dendrite targeting via a combinatorial cell-surface code. Neuron.

Xu, S., Xiao, Q., Cosmanescu, F., Sergeeva, A. P., Yoo, J., Lin, Y., Katsamba, P. S., Ahlsen, G., Kaufman, J., Linaval, N. T., et al. (2018). Interactions between the Ig-Superfamily Proteins DIP-α and Dpr6/10 Regulate Assembly of Neural Circuits. Neuron 100, 1369–1384.e6.

Xu, C., Theisen, E., Maloney, R., Peng, J., Santiago, I., Yapp, C., Werkhoven, Z., Rumbaut, E., Shum, B., Tarnogorska, D., et al. (2019). Control of Synaptic Specificity by Establishing a Relative Preference for Synaptic Partners. Neuron 103, 865–877.e7.

Xu, S., Sergeeva, A. P., Katsamba, P. S., Mannepalli, S., Bahna, F., Bimela, J., Zipursky, S. L., Shapiro, L., Honig, B. and Zinn, K. (2022). Affinity requirements for control of synaptic targeting and neuronal cell survival by heterophilic IgSF cell adhesion molecules. Cell Reports 39, 110618.

Yamagata, M. and Sanes, J. R. (2019). Expression and Roles of the Immunoglobulin Superfamily Recognition Molecule Sidekick1 in Mouse Retina. Front Mol Neurosci 11, 485.

Zhan, X.-L., Clemens, J. C., Neves, G., Hattori, D., Flanagan, J. J., Hummel, T., Vasconcelos, M. L., Chess, A. and Zipursky, S. L. (2004). Analysis of Dscam Diversity in Regulating Axon Guidance in Drosophila Mushroom Bodies. Neuron 43, 673–686.

